# The life span of steps in the enzyme-catalyzed reaction, its implications, and matters of general interest

**DOI:** 10.1101/2023.01.08.523157

**Authors:** Ikechukwu Iloh Udema

## Abstract

There is increasing recognition for steps and their life span (LS) in the enzyme-catalyzed reaction pathway; the need for rate constants, activation parameters, and transition states (TS) has assumed preeminence in the literature. The determination of the life span of complexes, enzyme-product (EP) complexes in particular, is not a regular feature in most studies. The study aims to show that there is LS for EP and TS destined for irreversible product formation and release. The cognate objectives are to: 1) derive the equation for the graphical determination of the first-order rate constant (FORC) of EP dissociation into E and P as well as the FORC of ES dissociation to free enzyme, E, and free substrate, S; 2) derive a FORC equation for the TS to EP conversion; 3) calculate the duration; and 4) calculate the FORC for the TS to EP conversion. Using mesophilic alpha-amylase and the Bernfeld method of assay, the velocities of catalysis were used to generate first the Michaelian parameters and then the rate constants. The LS of the TS and the FORC for the conversion of the enzyme-substrate (ES) complex to TS are 4.321 exp. (−6) min and 2.314 exp. (+5)/min, respectively; the LS of the TS destined for backward conversion (deactivation) to ES and its FORC are 7.525 exp. (−7) min and 1.387 exp. (+6)/min, respectively; the LS of the EP and its FORC for dissociation to E and P are 1.33 exp. (−5) and 7.52 exp. (+4)/min respectively. In conclusion, there are always steps in an enzyme-catalyzed reaction in the catalytic cycle; the LS of each step is shorter than the total LS of deactivation and dissociation processes; this is applicable to forward reactions. Thermodynamic activation parameters must account for all FORC in accordance with the additivity principle.

## 1. INTRODUCTION

Enzymes are well-known biological catalysts in that they are available in any phylum. Their mystery lies in their capacity to speed up biochemical reactions on a microsecond time scale or much less than that. The rates and cognate rate constants of their catalytic activity are of general interest to scientists in theory and practice. They are used to catalyze a wide range of commercially important processes, such as the production of sweetening agents and the modification of antibiotics; they are used in washing powders and various cleaning products; and they play a key role in analytical devices and assays that have clinical, forensic, and environmental applications. [1]. Other none-protein enzymes, such as ribozymes known in nature for gene expression and a man-made enzyme called abzymes, have both industrial and therapeutic applications. [1]. Wilhelm Kühne, a German physiologist, coined the term “enzyme” in 1878 [1], which is derived from the Greek words “en,” meaning “within,” and “zume,” meaning “yeast.” [1]. The term, enzyme reaction pathways have been of interest many years ago in the light of the fact there are steps before product is formed and released [2]. There has always been effort directed at the formulation of rate equations for enzyme-catalyzed reactions. The parameters that have attracted much attention are the Michaelian parameters, the *K*_M_ and *k*_cat_, and, more recently, the *k*_cat_/*K*_M_ [3, 4]. The latter has been criticized as being inadequate to describe the efficiency of a biocatalyst, particularly when employed as a criterion for selecting between enzyme variants for industrial purposes [5].

As stated earlier, so much interest lies in the quantification of Michaelian parameters, and in a preprint [6], it was opined that it may take extraordinary mathematics to be able to quantify the duration of the transformation of the transition-state (TS) taken as the activated enzyme-substrate complex (E^#^S^#^) to the enzyme-product (EP) complex; the E^#^S^#^ or TS is said to be in equilibrium with the ES [7]. Because the apparent catalytic rate constant is a function of activation energy, the goal of this study is to determine the duration (the life span) of physicochemical events in the enzyme-catalyzed reaction pathway and implications that are “en” the enzyme’s active-site, as well as to broaden the time-zones to include the duration of the TS→EP transition. To accomplish these aims, the research has the following objectives: 1) derive the equation for the graphical determination of the first-order rate constant (FORC) of EP dissociation into E and P as well as the FORC of ES dissociation to free enzyme, E, and free substrate, S; 2) derive a FORC equation for the TS→EP conversion; 3) calculate the duration of TS→EP conversion; and 4) calculate the FORC for the TS→EP conversion.

## 2. THEORY

The starting point in this rederivation is to state the background ordinary differential equation similar to the observation elsewhere [8] and to advise that phrase “such as after some algebra” is avoided to address all levels of intellectual achievement-not just high-rank scholars neck-deep in mathematical biosciences.

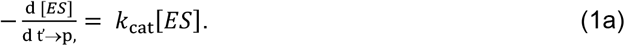

However, one may realize that [*ES*] = [*E* _f_] [*S*_f_]/*K*_M_ and substitution into Eq. (1a) gives:

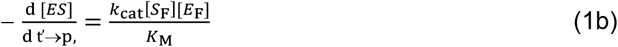

where ES, [*S*_F_], t’_→p_, *k*_→P_, and *K*_M_ are enzyme-substrate complex, concentration of free substrate, time, pseudo-first order rate constant for utilization of the substrate leading to product formation, and Michaelis-Menten (MM) constant respectively. Equation (1a) is set out as it is, because the dissociation of ES into free enzyme (E) and product (P) is considered. In such situation, [*ES*] is decreasing while, [*E*_F_] is increasing. However, this constitutes a simplification because, as indicated elsewhere [9] the activated complex needs to be accounted for as follows:

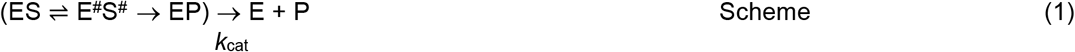

In scheme 1, E^#^S^#^, EP, *k*_cat_, and P are the activated enzyme-substrate complex, enzyme-product complex, catalytic rate, and product respectively. Further analysis of scheme 1 for the purpose of derivation comes up shortly. Given that, [*E*_F_] is equal to [*E*_T_]−[*ES*], and [*S*_F_] is equal to exp. (−*k*_→P_ t’_→p_,) [*S*_T_], where [*S*_T_] is the total substrate concentration in time, t’_→p_, equal to zero, then, Eq. (1) is restated as:

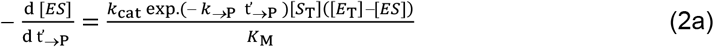

Rearrangement for the purpose of integration gives:

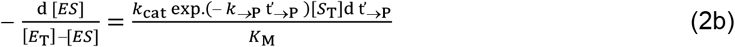

Upon integration of Eq. (2b), one obtains:

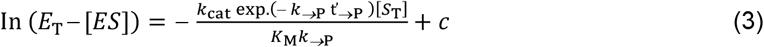

where *c* is an arbitrary constant. Since it is the extant ES that proceeds to E^#^S^#^ and ultimately to P and the free enzyme E, there is an initial build-up of ES. The time t’_→p_ is that time between the accumulation of ES, catalytic action, which entails activation, bond breaking and making, *etc*., and the transition to EP [9] before dissociation into free E and P. In other words, it is the period preceding the transformation of *E*^#^*S*^#^ into EP. This opens the door to determining the duration of the process, E^#^S^#^→ EP. The existence of ES and E^#^S^#^ occurs before E^#^S^#^ transforms into EP. The EP cannot reverse to E^#^S^#^ due to thermodynamic considerations, as per its infeasibility [9].

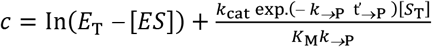

At t’_→p_ equal to zero,

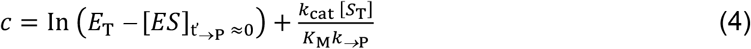

[*ES*]_t′ ≈0_ > 0: However, after time, t’→p, the time required to convert the substrate into a product and a free enzyme, the [*ES*]_t’_ = 0. Substitute Eq. (4) into Eq. (3) to give:

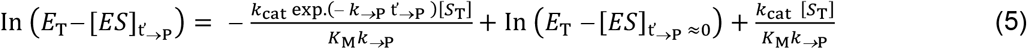

Rearrangement of Eq. (5) gives:

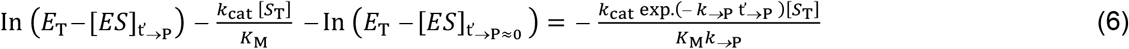

Further rearrangement gives:

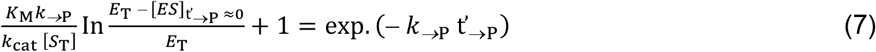

Taking the natural log gives:

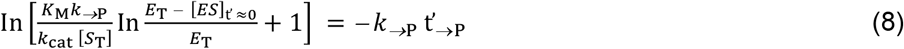

Equation (8) may give rate, *i*.*e*. the reciprocal of t’_→P_ similar to *k*_cat_. Thus, Eq. (8) may require further modification to address specifically the dissociation of EP to free enzyme and product-EP→E + P. This is to be addressed shortly.

Equation (7) is as such because, as stated earlier, when t’_→p_, is > 0 (i.e., the duration of the process, (E^#^S^#^→EP)→E + P), the [ES] leading to E^#^S^#^ and eventually to E and P does not exist; this is not to imply that elsewhere in the reaction mixture, there are no new formations of ES. This is a case-by-case situation. A plot of the left-hand side (LHS) versus *k*_→P_ gives a slope equal to t’_→p_. The derivation of *k*_→P_ has been described in the literature [10]. There should be a pseudo-first-order rate constant for cases in which the probability that ES cannot progress to product formation—the case in which E^#^S^#^ is deactivated—is high. For such a case, the equation equivalent to the equation for *k*_→P_ is stated as follows:

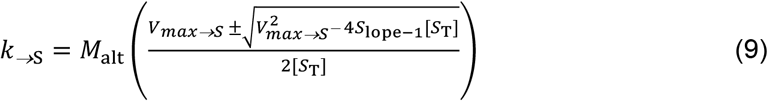

where *M*_alt_ and *V*_max→S_ are the molar mass of maltose and the maximum velocity of the process given as: E^#^S^#^ → ES→E + S; *V*_max→S_ = *k*_-1_[*E*_T_] where *k*_-1_ is the first-order rated constant for the process given as: E^#^S^#^ → ES→E + S. The slope (S_lope-1_) is obtained by plotting *v* versus [*S*_T_]/(*V*_max→S_ − *v*); *v* is the experimental variable-the velocity of catalysis. There is no hard and fast rule as to the method for the determination of *k*_→P_ and *k*_→S;_ otherwise, there are other methods in the literature [11]; researchers only exercise their discretionary right to choose conveniently without making any of the options sacrosanct or a definitive model.

After putting in place the necessary equations for calculating *k*_→S_, Eq. (8) must be restated for the process duration (t’_→S_), ES → S + E, as follows:

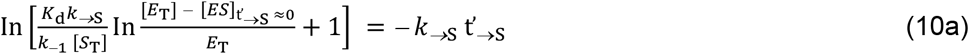

where *K*_d_ is the dissociation constant of ES given as *k*_−1_/*k*_1_. A plot of LHS versus *k*_→S_ gives a slope equal to t’_→S_. In the light of Eq. (10a), its equivalent equation for the dissociation of EP to free E and P should be:

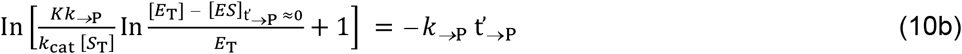

where *K* is the Van Slyke–Cullen [12] constant

However, in line with the aim of this research, the earlier assumption [6] that it may be impossible, as of then, to determine the duration of the process, *E*^#^*S*^#^→EP is hereby reversed in favor of the possibility of doing so as follows: First, the reciprocal of the rate, 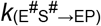, in the equation below gives the duration or life span of the process, (E^#^S^#^→EP).

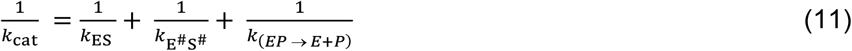

where 1/*k*_ES_, 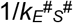, and 1/*k*_(EP→E+P)_ are the durations of the processes, ES formation, activation of ES, and dissociation of EP to free enzyme, E and P respectively.

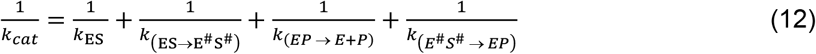

Making *k*_cat_ subject of the formula requires that:

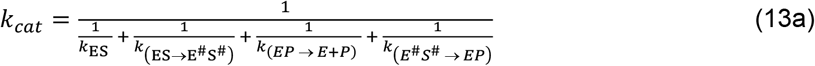

In terms of the durations expressed in the symbol of time (life span) used in this study:

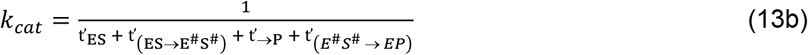

In this research t’_(EP→E+P)_ is equivalent to t’_→P_ (Eq. (8)). From Eq. (12) is given the following:

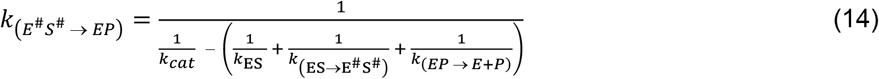

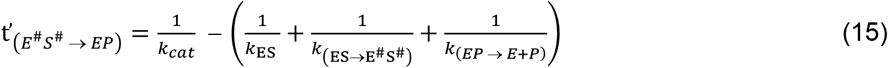

Equations (14) and (15) are an expression of the realization of the objectives of this research. However, there must be information about 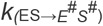 in order to calculate the dependent parameters in Eqs (14) and (15). Eq. (13b) gives the equation for determining k_(EP→E#S#)_ → (*k*_+2_ for short) as follows:

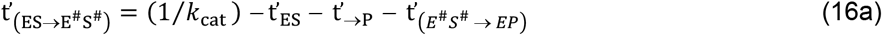

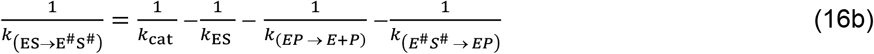

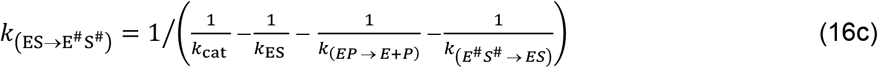

The determination of *k*_ES_ is as described in the literature [13].

Meanwhile, it needs to be made clear that the size of the *V*_max_ depends not just on the concentration of the enzyme and the saturating concentration of the substrate, but also on the concentration of the stable transition state complex (E^#^S^#^) that progresses to EP after the chemistry of the process is concluded. As a result, if the magnitude of the stable E^#^S^#^ and thus EP is less than the minimum threshold, the *V*_max_ cannot be reached. This is not to say that the TS (or E^#^S^#^ as a reminder) and EP must, with time, eventually be equal [*E*_T_]: The trend in the magnitude of the former with time should indicate the probability that *V*_max_ may be attained. Hence, just as the *k*_cat_ is the same for the same enzyme in a defined condition, regardless of its concentration, as long as the [*S*_T_] is much greater than the [*E*_T_], the first-order rate constant for the conversion of E^#^S^#^ to EP—the second aspect of the chemistry as a strict function of the enzyme—is also the same for the same enzyme regardless of its concentration.

Under the same assay conditions, the enzyme exhibits the same mechanism of catalysis within the same time period, yielding the same rate constant. The magnitude of ES and TS depends on the concentration of E and S. The first aspect of the chemistry is the formation of TS. This aspect can be aborted for reasons ranging from thermal energy-orchestrated perturbation leading to catalytic disorientation of the vulnerable groups of substrate and the functional groups aiding catalysis and located close to the active site of the enzyme. For carbohydrate as a substrate, the carbohydrate binding modules (CBM) may be relevant in this regard. The occurrence of CBN in the N-terminal region of the Laforin gene, perhaps of the kind “coding for a protein-phosphatase involved in glycogen metabolism,” has been recognized [14]. The opinion in the literature is that a single mutation in CBN, reducing its carbohydrate binding capacity, causes Lafora disease [14].

It is documented that there is a dynamic equilibrium between ground-state ES and its TS counterpart [7]. Therefore, there are clearly two first-order rate constants, “forward” and “backward,” 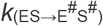 and 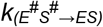, respectively. These first-order rate constants 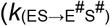 and 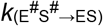 are now denoted by *k*_2+_ and *k*_-2_. The former symbols represent chemistry, specifically activation and deactivation processes. When *k*_-1_ is much greater than *k*_cat_, it means that a preponderance of the process, E^#^S^#^→ES, is the case; the converse is the case if *k*_cat_ is greater than *k*_-1_. [15]. From the chemistry of the process in which the enzyme assumes the leading role, the conversion of ES to TS is regarded as occurring at the same rate for the same enzyme and substrate under clearly defined and specific assay conditions. This assertion is also applicable to the conversion of TS to EP, the second aspect of the chemistry. The magnitude of the rate of deactivation (E^#^S^#^→ES,) affects the magnitude of *k*_−1_, and the magnitude of the rate of activation (TS formation-ES→E^#^S^#^) affects the magnitude of *k*_cat_. Thus, it makes sense to state that the ratio of *k*_−1_ to *k*_cat_ may be equal to the ratio of *k*_−2_ to *k*_+2_. With such an equation, one can state the equation for *k*_+2_ as follows:

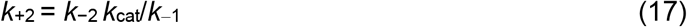

Next one can derive composite equations for *K*_M_ in terms of *k*_−1_ and *k*_cat_ as follows as corollaries. Firstly, from 1/*k*_−1_ = 1/*k*_11_+1/*k*_−2_ where *k*_−2_ is ≡ *k*_(ES→E+S)_ and *k*_11_ is ≡ *k*_(ES→E+S)_, is given:

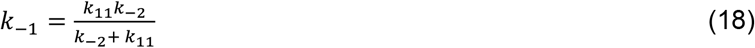

Meanwhile, the Michaelian equation for the *K*_M_ gives two equations: a) *k*_−1_ = *K*_M_*k*_1_ − *k*_cat_ and b) *k*_cat_ = *K*_M_*k*_1_ − *k*_−1_; thus, with Eq. (a), the equation of *K*_M_ in terms of *k*_cat_ and the equation of *k*_−1_ as a function of all first-order rate constants for the processes leading to the release of free S and free E is given as:

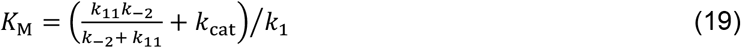

Meanwhile, Eq. (12), is restated as:

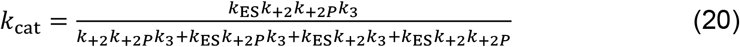

where 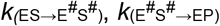, and *k*_(EP→E+P)_ are redesignated as ≡ *k*_+2,_ *k*_+2P_, and *k*_3_ respectively.

The *K*_M_ in terms of *k*_−1_ and the equation of *k*_cat_ in terms of all the first-order rate constants for the processes leading to the release of P and free enzyme are then, with Eq. (b), given as:

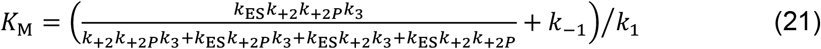

After substituting Eq. (b) into Eq. (20) and rearrangement it becomes:

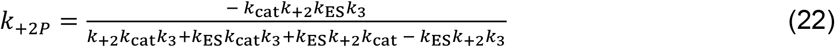

On matters of general interest, one can begin discussion by reminding the scientific community that biochemistry, biophysics, computational biology (or biochemistry), and chemistry are quantitative sciences. Therefore, it is often the case that the Arrhenius activation energy is graphically determined, such that not less than three different temperatures may be needed for such a plot while the Gibbs free energy of activation is calculated for each temperature as noted elsewhere [10]; unfortunately, the Eyring-Polanyi equation and the Arrhenius equation seem to be used interchangeably [7]. Previous studies have entered into the argument against this approach even if there was, as in most respected top-class journals, the improper use of *k*_cat_ as a rate constant for only a step in the catalytic cycle [10]. For instance [10],

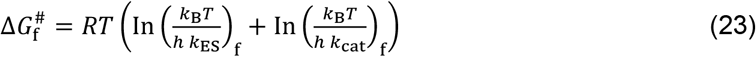

where 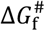, *R, T, k*_B_, and *h* are Gibbs free energy of activation in the forward (f), direction, universal gas constant, thermodynamic temperature, and Boltzmann constant respectively. Although Eq. (23) is conceptually similar to the equation to be derived in this research, the presence of *k*_cat_ is inappropriate in the light of the current conceptual predisposition in this research. Also of concern is the following [7] definition of the Eyring-Polanyi equation:

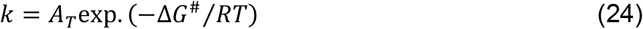

where *k* (a general designation for first-order rate constant) is a first-order rate constant. The equation is not very different from

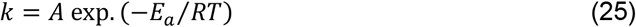

where *E*_a_ and *A* are the Arrhenius activation energy and pre-exponential factor respectively; according to Kohen [7], the function *A* (intended to mean *A*_T_) includes all the pre-exponential terms, such as the transmission coefficient (κ), friction factors, re-crossing events, *etc*. The implication is that *A* is = κ*k*_B_*T*/*h*. Yet the older Arrhenius equation requires that In*k* is plotted versus 1/*T*; but the Eyring-Polanyi equation is given as:

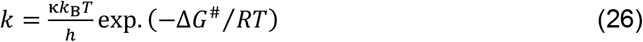

Plotting Ink versus 1/*T* from equation (26) should yield different results than equations (24) and (25); this is based on the fact that the Arrhenius equation has nothing to do with κ or any other factor. But this remains speculative and re-emphasizes the issue of controversy surrounding Arrhenius (and recent variants) and Eyring-Polanyi equations [10]. It seems the only thermodynamic activation parameters that should be graphically determined are the Arrhenius activation energy and its cognate enthalpy of activation, which is obtained by subtracting *RT* from *E*_a_. Otherwise, Eqs (27) and (24/25) need to represent different parameters in order to settle the controversy.

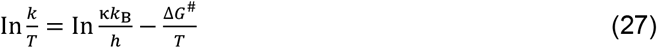

Regardless of the controversy, Arrhenius activation is recognized at all levels as the minimum amount of energy required by the reactants to initiate the reaction; this should determine the kinetic aspect in the formation of encounter-complexes before enzyme-substrate complex formation. The newer entrant, the Gibbs free energy of activation, is the difference between the free energy of the reactive reactants and the free energy of all reactants [16]. A clearer definition is that it is the free energy difference between the reactant state (RS) and the corresponding transition state (www.sciencedirect.com). The adoption of the additivity principle in the computation of Gibbs free energy of activation resulted from this study, which should serve as an improvement to Eq. (23). Keeping constant (and assuming = 1), a general expression for the summed Gibbs free energy of activation is as follows:

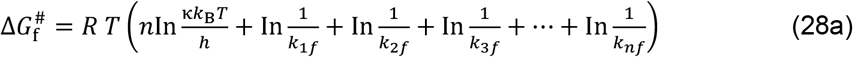

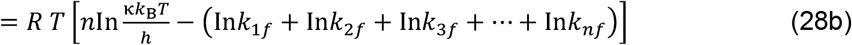

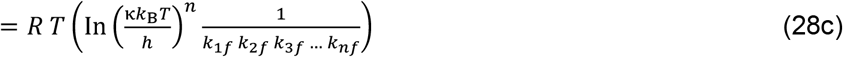

In the light of this study, the specific equation is given as:

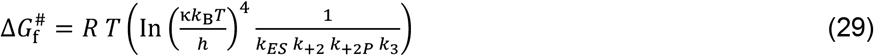

The backward equivalent of Eq. (28c) is:

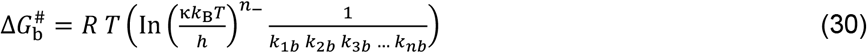

For the purpose of this investigation, the equation is:

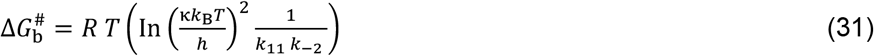

As a matter of common sense, if 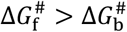, the energy “barrier height” against the formation of product is high and the rate of product formation may be low and *vice versa*.

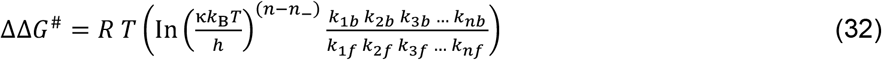

For the purpose of this study the equation is:

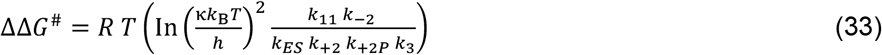

*n*−*n*_−_ is = 2. For the graphical determination of the free energy of activation the following equation may be required.

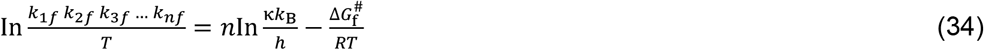

The equation for the backward equivalent is:

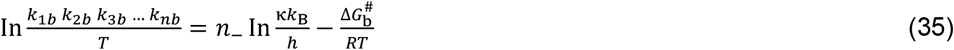

On grounds of sincerity and “intellectual humility” and for the sake of “pedagogical demands”, Eqs (34) and (35) need to be re-examined: Because 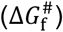 and 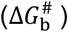 are all generally equal to Δ*H*^#^ −TΔ*S*^#^, substituting the latter into, say, any of the equations yields:

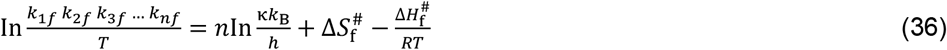

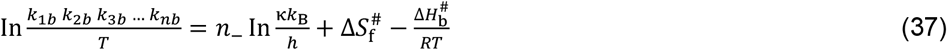

where *S* and *H* denote the entropy and the enthalpy respectively. Eqs (36) and (37) make no mention of innovation, but it should be noted that the same slope governs the value of the enthalpy and free energy of activation. To put it another way, the same slope is = Δ*G*^#^/R = Δ*H*^#^/R**!** Without the “International Space Center Observatory,” one can clearly observe that with the original Arrhenius equation (not to the exclusion of its variants), the following equations can be given:

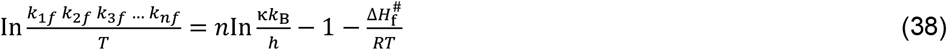

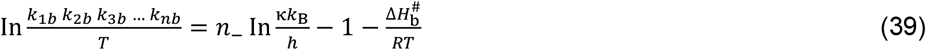

Therefore, given any value of *n* the intercept is clearly, *n*In(κ*k*_B_/*h*) – 1; if a unimolecular reaction is inferred *E*_a_ should be = Δ*H*^#^ + *RT*: Here again, the latter can be substituted into original Arrhenius equation to give:

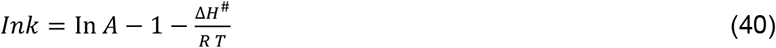

Equations (38), (39), and (40) are similar. Yet, the *R* times the slope in Eq. (40) cannot give different values of the enthalpy of activation and Arrhenius activation energy. It may require high-ranking chemical physicists, physical chemists, *etc*. to come up with a clear-cut redefinition with a clear contrasting feature in future equations that can distinctly be applied to Arrhenius activation energy and thermodynamic activation energy. Either a plot gives first the enthalpy of activation, which can be added to *RT* to give *E*_*a*_, or the ΔS^#^ obtained by subtracting 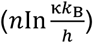 from the intercept (Eqs (36) and (37)) is added to the same enthalpy of activation to give the Gibbs free energy of activation. After all, the heat content of a reaction mixture is known as its enthalpy; it can be increased by the supply of external heat, which in most cases accelerates the velocity of reactions, including those catalyzed by enzymes, as long as the enzyme remains structurally and conformationally stable.

Activation energy is calculated, but heat content is instrumentally measured. If the minimum kinetic energy that reactants need to attain TS and commence chemical reactions is activation energy, and the former is a state function like Gibbs free energy of activation, then Arrhenius activation energy may be a state function; this can be redefined as the minimum amount of work needed to overcome the energy barrier against the attainment of TS: The Gibbs free energy of activation, being a measure of the potential energy barrier, subsumes the Arrhenius activation energy, which can also be regarded as a fraction of the Gibbs free energy of activation needed to start a reaction. The deductions that can be made are that the original Eyring-Polanyi equations or any other variant and the Arrhenius equations or any other variant may have been directly relevant to the graphical determination of enthalpy of activation. Speculative as this may be, the derivational analysis made so far cannot be speculative because a valid equation that is dimensionally accurate cannot reproduce falsity.

### 3.1. MATERIALS AND METHODS

#### 3.1.1 Materials

##### 3.1.1.1 Chemicals

The enzyme which was assayed is *Aspergillus oryzae* alpha-amylase (EC 3.2.1.1) and, insoluble potato starch, was the substrate; both were purchased from Sigma–Aldrich, USA. Tris 3, 5—di-nitro-salicylic acid, maltose, and sodium potassium tartrate tetrahydrate were purchased from Kem Light Laboratories in Mumbai, India. Hydrochloric acid, sodium hydroxide, and sodium chloride were purchased from BDH Chemical Ltd., Poole, England. Distilled water was purchased from the local market. The molar mass of the enzyme is approximately 52 k Da [17,]. Distilled water was purchased from the local market. As a word of caution, as advised in previous publication [18] readers of this paper should be aware that the use of the same enzyme in articles by the same author(s) is strictly due to budgetary constraints. In this research much lower concentrations of the substrate and the enzyme are explored to make a difference (see preparation of reagent and solution of enzymes below). The evaluation of new models remains a primary concern.

##### 3.1.1.2 Equipment

An electronic weighing machine was purchased from Wensar Weighing Scale Limited, and a 721/722 visible spectrophotometer was purchased from Spectrum Instruments, China; a *p*H meter was purchased from Hanna Instruments, Italy.

### 3.2 Methods

#### 3.2.1 Preparation of reagents and assay

The method of assaying the enzyme is Benfield’s method [19], with gelatinized potato starch as substrate, whose concentration range is 3–10 g/L. The reducing sugar produced upon hydrolysis of the substrate at room temperature (≈ 24 _°_C) using maltose as a standard was determined at 540 nm with an extinction coefficient approximately equal to 181 L/mol.cm. The duration of assay is 3 minutes. A mass concentration equal to 0.2 mg/L of *Aspergillus oryzae* alpha-amylase was prepared in a Tris-HCl buffer at *p*H 6.0; there were no special considerations in the choice of *p*H and temperature. The evaluation of new equations was the only overriding interest.

#### 3.2.2. The determination of rate constants

The pseudo-first-order rate constant for the utilization of gelatinized starch is determined as described in the literature [10], while the second-order rate constant for the formation of the enzyme-substrate (ES) complex and the duration of its formation are determined as described elsewhere [13]. The first-order rate constant, *k*_−1_, for the release of free E and free S in a two-step process is also determined as described in the literature [13]. The Lineweaver-Burk [20] plot was used for the determination of the *K*_M_ and *V*_max_. Equations (8) and (9) were for the calculation of *k*_+2_ (*i. e*., 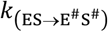) and *k*_-2_ (*i. e*.,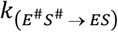) respectively. The determination of *k*_3_ (*i. e*., *k*_(*EP* → *E*+*P*)_) is as described in the literature [13], but a corrected version is rederived herein as shown in Eq. (8) and yet again in its specific form that addresses a specific catalytic event—the dissociation of EP to free E and P, where the reciprocal of the slope, t’_→P_, is equal to *k*_3_. The reciprocal of t’_→S_ (a slope) in Eq. (10) is the rate constant for the dissociation of ES to free E and free S in analogy to the dissociation of EP to free E and free P, as investigated in a manuscript awaiting evaluation [6].

Consequently, 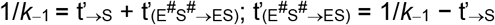.

### 3.3 Statistical analysis

Assays were conducted in triplicates. Micro-Soft Excel was used for the determination of the standard deviation (SD) for the arithmetic mean values.

## 4. RESULTS AND DISCUSSION

Preliminary data were generated by a Lineweaver-Burk plot [20] (Figure not shown). The data covers the Michaelian parameters, the *K*_M_ and *V*_*max*_, and the first-order rate constant for the formation and release of product, otherwise known as the catalytic rate, *k*_cat_ (Table 1). The *k*_cat_ value is needed for the calculated values of the independent variables, which are plotted versus the pseudo-first-order rate constant (*k*_→P_) for the utilization of substrate [13] for the determination of the duration of ES formation (Figure 1).

**Table 1:**
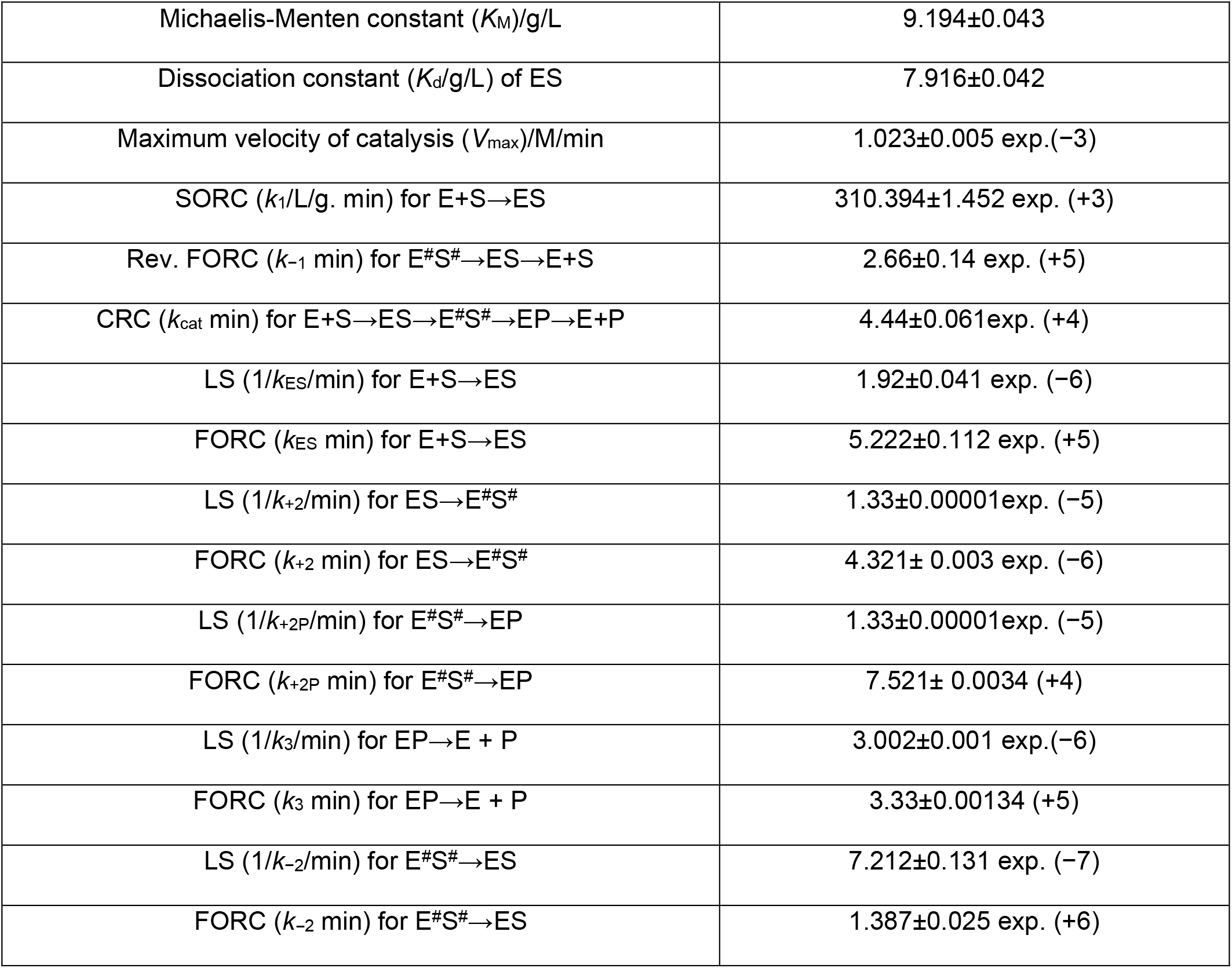
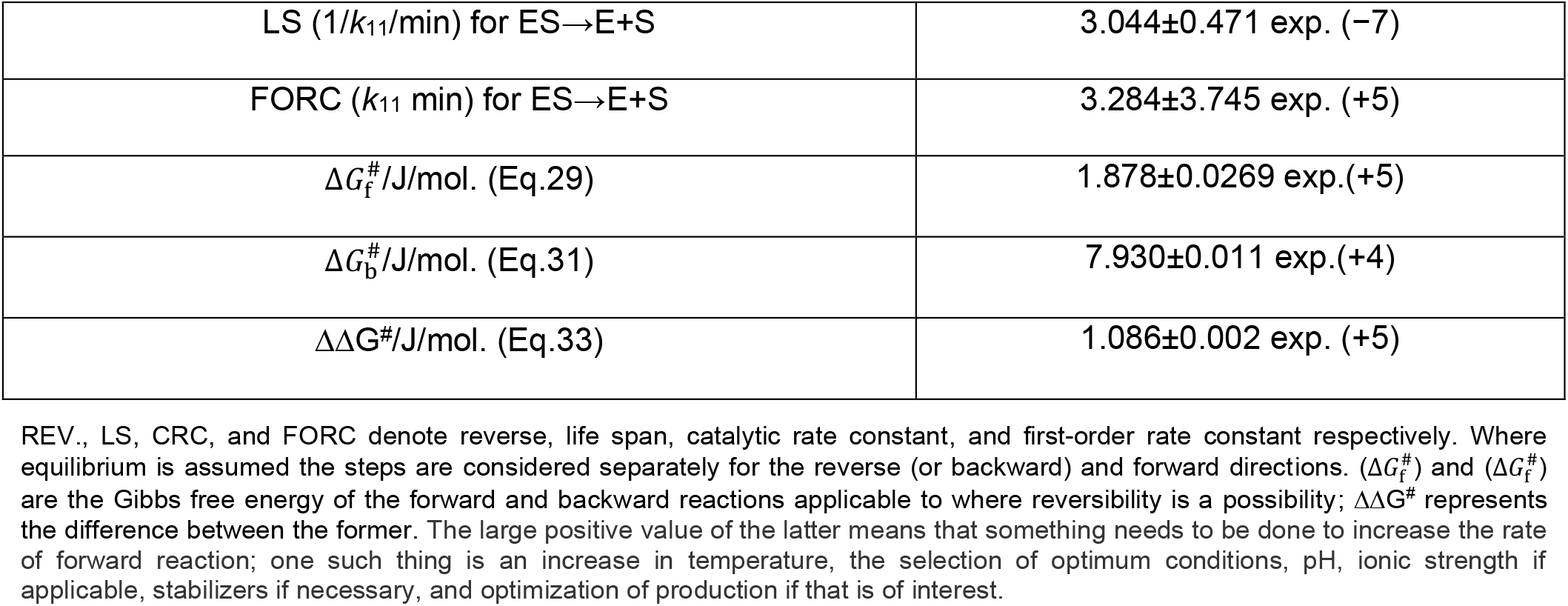
Kinetic parameters and activation energies obtained by calculation and graphical method

**Figure 1:**
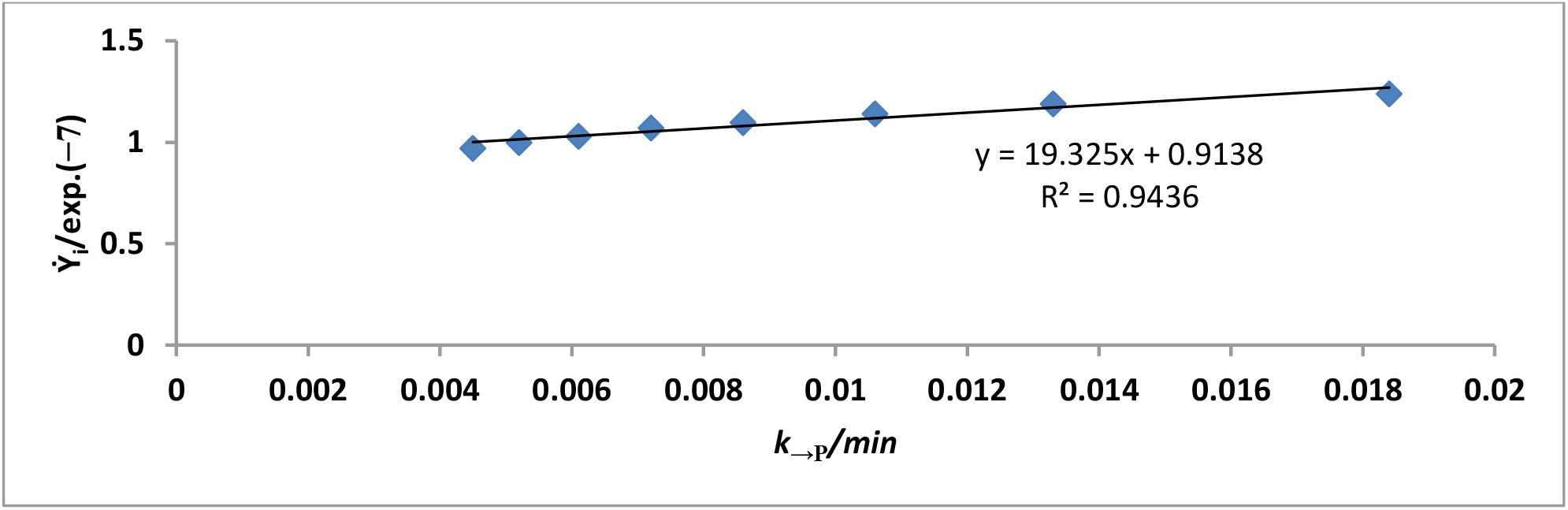
Determination of the duration of enzyme-substrate complex (ES) formation: *k→P* is the pseudo-first order rate constant for the utilization of the substrate and 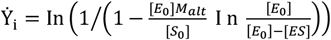. The slope of the graph is the duration of ES formation *i.e. t*_ES_.

In this study, one should keep in mind that the use of [*E*_T_] 0.2 mg/L with a substrate concentration range of 3–10 g, as opposed to 2.5 mg/L [13] and 1.667 mg/L [6] with a similar substrate concentration range (5–10 g/L), has implications in light of challenging criteria frequently discussed in the literature [21, 22]. In this study, one should keep in mind that the use of [*E*_T_] 0.2 mg/L with a substrate concentration range of 3–10 g, as opposed to 2.5 mg/L [9] and 1.667 mg/L [6] with a similar substrate concentration range (5–10 g/L), has implications in light of challenging criteria frequently discussed in the literature [21, 22]. In some cases, these assumptions have similar criteria, but none of them can justifiably repudiate the requirement that zero-order kinetics is a way to determine experimentally the maximum velocity of the catalytic action of an enzyme. The lower the concentration of the enzyme, the higher the *k*_cat_, though this is not a rule because the underlying reason is that a higher concentration of the enzyme requires a much higher concentration of substrate to achieve saturation; this cannot easily be defined. Thus, previous research found *k*_cat_ values of 1.566 exp. (+4)/min ([*E*_T_] = 4.8077 exp. (−8) M ≡ 2.5 mg/L) [9] and 2.108 exp. (+4)/min ([*E*_T_] = 3.205 exp. (−8) M ≡ 1.6667 mg/L) [6] that are significantly less than 4.437 exp. (+4)/min (3.846 exp. (−8) M ≡ 0.2 mg/L (Table 1)); these values correspond to [*S*_T_]/[*E*_T_] ratio of 2.39-4.8, 1.59-3.2, and 7.645-15.279 respectively. This analysis is in line with the advice that the assumptions under which an assay is carried out need to be indicated [23]. As an advocate (or better yet, as a “student”) of the Michaelis-Menten and Briggs-Haldane principles, this research is much closer to the criteria that justify sQSSA.

The use of *k*_cat_ is also applicable to the graphical determination of the zero-order second-order rate constant (*k*_1_) for the formation of ES. The generated independent variables calculated using *k*_cat_ as described in the literature [13] are plotted versus *k*_→P_ (Figure 2). The slope is the value of *k*_1_ (Table 1). The name zero-order second-order rate constant implies that *K*_M_, an outcome of a mixed-order zone, and *V*_max_ (or its alternative, *V*_max_/[*E*_0_] = *k*_cat_), are part of the equation of the Michaelis-Menten constant given as: *K*_M_ = (*k*_-1_+*k*_cat_)/*k*_1_; besides, a method for graphical determination of *k*_1_(=(*k*_-1_+*k*_cat_)/*K*_M_) is known [as derivable]. The value of *k*_1_ obtained in this study (Table 1) is 11.48 exp. (+6) L/mol. min. (33.567 exp. (+3) L/g. min.), which is higher than 4.754 exp. (+6) L/mol. min. (13.901 exp. (+3) L/g. min.) reported in the literature for the same enzyme (2.5 mg/L) [9]. Also, the values of *K*_d_, *K*_M_, and *k*_1_ (17.61 g/L, 18.66 g/L, and 2.447 exp. (+5)/min, respectively) in the literature [9] differ from the values obtained in this study (≈7.92 g/L, 9.18 g/L, and ≈2.66 exp. (+5)/min, respectively) (Table 1).

**Figure 2:**
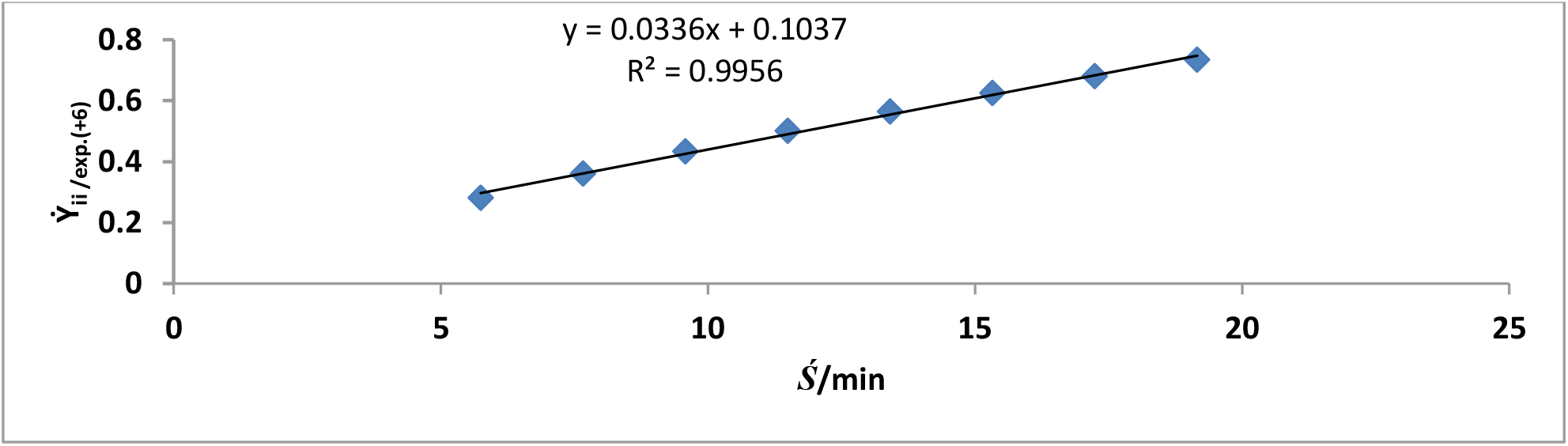
Determination of asymptote second-order rate constant for the formation ES. 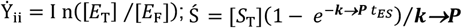. The slope of the graph is = (*k*_−1_+*k*_cat_)/*K*_M_.

In a previous study, Eq. (8), the rederived equation was used to determine graphically the value of t’_→P_; but such an effort may yield a value whose reciprocal is similar to *k*_cat_. Therefore, Eq. (10b) addresses a specific catalytic event such as the dissociation of EP to free E and P. Generated independent variables where the Van Slyke-Cullen constant is relevant [12] were plotted versus the corresponding pseudo-first-order rate constant (***k***_**→P**_) for the effective utilization of the substrate (Figure 3). The plot gives a slope, the equivalent of the life span (Table 1) of EP. The obtained EP life span and first order rate constant for the release of P and free enzyme are 3.00 exp. (6) min and 3.33 exp. (+5)/min (Table 1), which are less than the values (3.17 exp. (6) min and 3.155 exp. (+5)) for a higher concentration of the same enzyme in a preprint server [6]. It must be revealed for building confidence that the equation (Eq. (10b)) used in this research is a modification of Eq. (8) used elsewhere [6].

**Figure 3:**
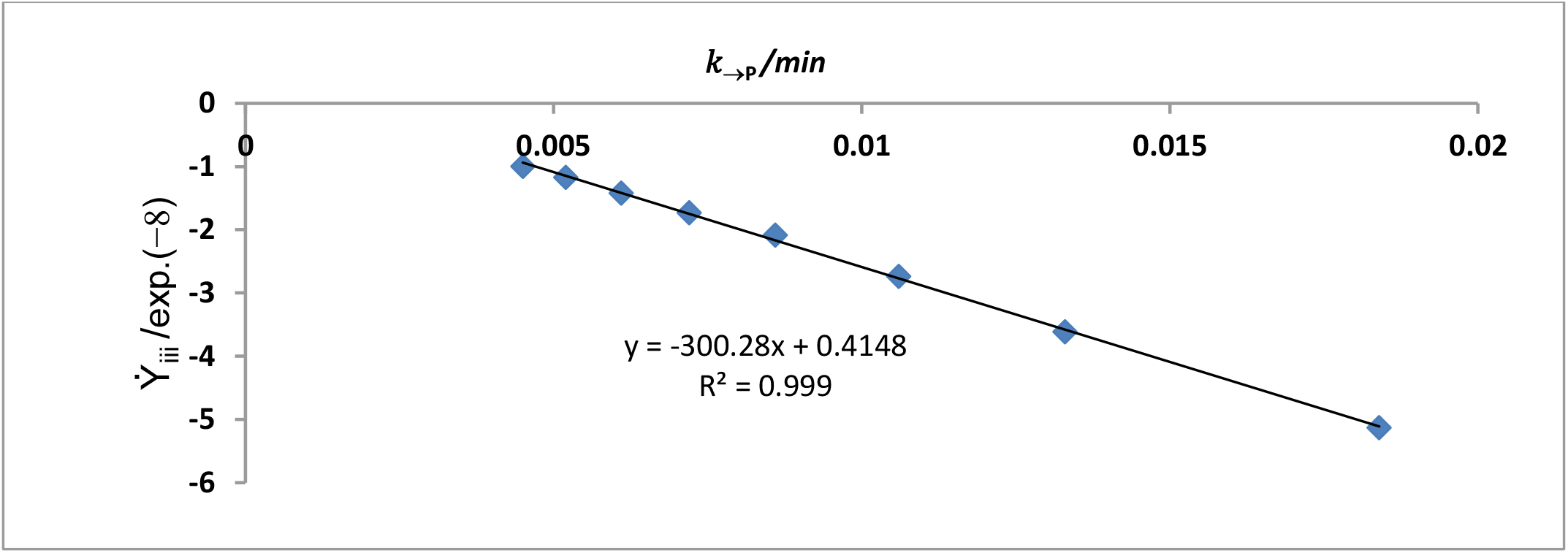
The determination of the life span or the duration for the dissociation of EP to free enzyme and product. 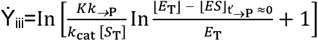. The slope of the graph is = the life span of EP; *k*_(→P)_ is the pseudo-first-order constant for the substrate destined to the amylolytic action of the enzyme.

As stated earlier, the adoption of the “zero-order” dissociation constant given as *k*_−1_/*k*_1_ reasonably necessitated the need to be more specific in the approach for the determination of the life span of the EP. Otherwise, there should be no basis for a matched comparison of the events that lead to the ES dissociation into free enzyme and free substrate and the EP dissociation into free enzyme and product. Here, in this research, any substrate that is destined to remain complexed with the enzyme, leading to steady TS, is assumed to differ in its pseudo-first-order rate constant for its utilization. The substrate molecule, which dissociates from its complex following TS deactivation, is regarded as having a pseudo-first order rate constant for its utilization, as opposed to its counterpart, which is certain to be hydrolyzed into product due to its stability. The values (not shown) of *k*_→S_ and *k*_→P_ were calculated as described with Eq. (9) and in the literature, respectively [10, 24].

The slope of a plot of the calculated independent variables versus *k*_→S_ (Figure 4) corresponds to the life span (3.04 exp. (−6) min) of unstable ES destined to dissociation to free S and free E; the corresponding first-order rate constant is 3.284 exp. (+5)/min. This rate is much less than the value (5.375 exp. (+6)/min) reported in a preprint [6]. The *k*_EP→P_ values for the dissociation of EP to E and P are slightly greater than the *k*_ES→E +S_ values for the dissociation of ES to free S and free E.

**Figure 4:**
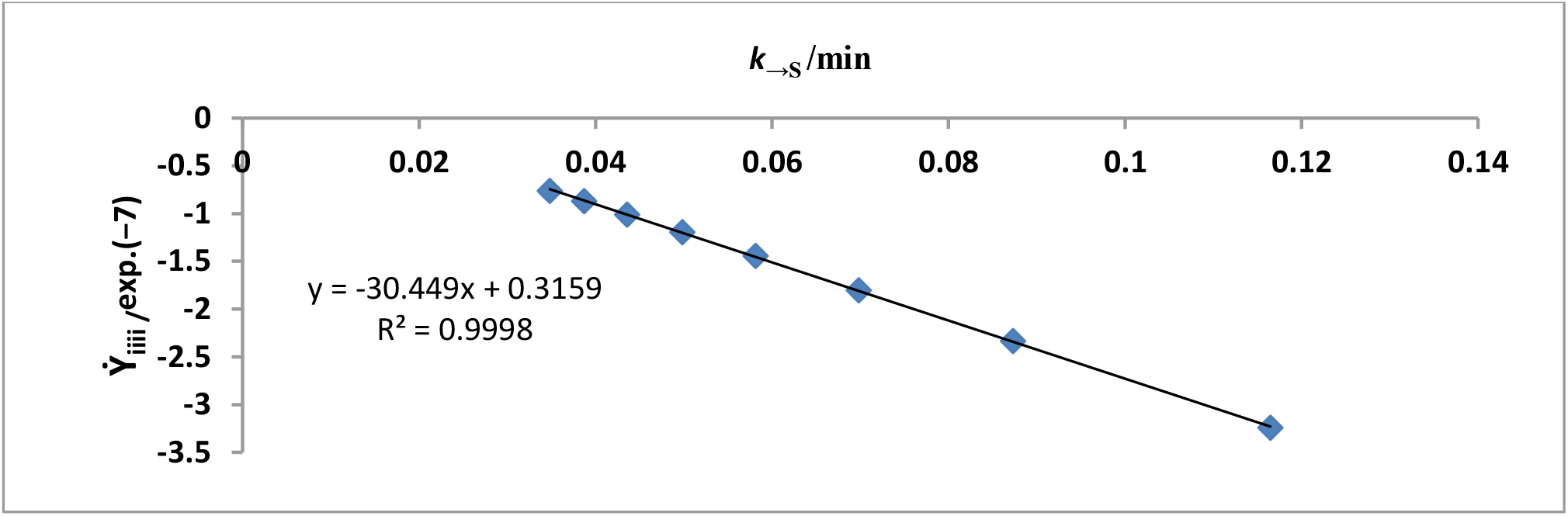
The determination of the life span or the duration for the dissociation of ES to free enzyme and substrate. 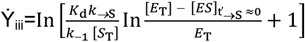. The slope of the graph is = the life span of ES; *k*_(→S)_ is the pseudo-first-order constant for the substrate destined to dissociate to free enzyme and free substrate.

The main goal of this study is, for the first time, to derive a first-order rate constant equation for the process, E^#^S^#^→EP, a process that is considered irreversible if, in particular, the enzyme is neither a synthase nor a synthetase. A pre-print paper [6] showed how the first-order rate constants for the processes ES→E^#^S^#^ and (E^#^S^#^⇌EP)→E+P can be determined; however, the process in parenthesis was considered not amenable to a derivable equation for the determination of its first-order rate constant and its life span. However, in this study, the first-order rate constant equations for ES→E^#^S^#^ and E^#^S^#^→E+P are given as Equations (17) and (22). The first-order rate constants arising from the two equations are 2.314 exp. (+5)/min (the corresponding life span is = 4.321 exp. (−6) min.), which is higher than the preprint’s [6] reported value, and 7.521 exp. (+4)/min (the corresponding life span is = 1.33 exp. (−5) min), respectively. Most important is the calculated value of the first-order rate for the process, E^#^S^#^⇌EP; this was found to be 7.521 exp. (+4)/min. This is ≈ 1.7-fold higher than *k*_cat_. With a corresponding life span of ≈ 1.33 exp. (−5) min, it goes to reaffirm that the life span of each step in the catalytic cycle is less than the duration (2.254 exp. (−5) min) of the catalytic cycle in the forward direction, to be specific; this has become a topical issue in recent time in the literature [4, 25].

In this study, 1/*k*_ES_ + 1/*k*_+2P_ + 1/*k*_+2_ + 1/*k*_3_ (≡ t’_ES_ + t’_+2P_ + t’_+2_ + t’_3_) is ≈ 2.253496 exp. (−5) min. The difference (≈0.0224 % of 1/*k*_cat_) is as a result of imperfection in measurement and experimental error. The duration (1/*k*_−1_ ≈ 3.766 exp. (**−**6) min) of the backward reaction is very similar to the sum of the life span (3.76581 exp. (−6) min) of the steps; the difference is ≈ 0.00658 % of 1/*k*_−1_. This result clearly supports the view in the literature that “the rare event of the barrier crossing toward products is likely to be on faster time scales than the first and following events” [7] in the catalytic cycle.

One can comment that despite the variety of ways in which the life span of each step in the catalytic cycle is determined, in both forward and backward directions, there is excellent agreement in the arithmetic relationship between the life span of each cycle and the sum of the life spans of individual steps; note that the life spans of the EP and ES (the latter undergoes dissociation in the backward direction) were graphically determined. Indeed, *k*_−1_ and *k*_cat_ were determined by graphical and calculational methods. These may imply an expression of the validity of the current models of rate constant equations and underlying methods. Nonetheless, there is an increasing preference for specificity constant [3], despite what one might consider a “reductionist approach” in the determination of the life span of specific steps in the catalytic cycle. One may wish to add that the reciprocal variant of the direct linear plot [26] can just certify the test for a unified or direct measurement of the specificity constant.

Here lies the importance of rate constants, the first-order rate constants in particular, where consideration is given to kinetic and thermodynamic measurements; kinetic measurements, taking into account first-order rate constants, can serve to define thermodynamic parameters governing enzyme specificity [27]. The Gibbs free energy of activation and the apparent free energy are important in this regard. The chemistry of a catalyzed reaction is a function of the catalyst, the enzyme, in this study. The magnitude of first-order rate constants is an expression of the conformational state of the enzyme. Failure of an enzyme to change from a ground state (GS) to a transition state (TS) means that the enzyme cannot assume the catalytic conformational state needed to lower the energy barrier. This is in line with the view that some enzymes (not to be too generalistic) undergo mutual “induced-fit” shifting of the conformational ensemble upon binding; then, via thermal search of the conformational space, they proceed toward the reaction’s transition state (TS) [7]. This issue has been of interest to researchers [7]; this study produced various first-order rate constants that can be explored for the determination of Arrhenius activation energies or Gibbs free energies of activation in a reductionist fashion that can be summated for each type of activation.

As Table 1 shows, the sum of the Gibbs free energies of activation in the forward direction is ≈188 kJ/mol., while the value for the backward reaction is ≈79 kJ/mol. The difference between the two is ≈108.56 kJ/mol. The large positive value is a reflection of the high energy barrier that stands against the progress of the chemistry to the TS formation. According to Kohen [7] it appears that enzymatic systems did not effectively stiffen restricting motions orthogonal to the chemical coordinate, which would have otherwise enabled the selective occurrence of dynamics along the reaction coordinate. Doing so would have meant that the life span of the EP should have been much shorter than observed (Table 1). Herein lies the important implication of the life span of each step in the reaction pathway in both the forward and backward directions: if steps in the forward direction have a longer life span as opposed to a very transient duration or higher first-order constants, the amount of product formed should be less than the optimum values expected. The desire for the optimization of industrial production could be hindered.

The desire of doctors and pharmaceutical companies is to produce drugs that can control metabolic disorders. In this regard, a longer life span of chemical species is desirable; this requires partial but significant inhibition of target enzymes using prescribed medication for the control of diabetes for instance. There is a proposition that thermodynamics and kinetics that characterize an enzyme-catalyzed reaction can best be studied and analyzed given information about the rate constants that may be a reflection of the structural basis of the observed conformational transitions [27]. Such information may enhance the design of drugs for the treatment of disease [27]. Therefore, this study has provided models-the equations-that can be used to evaluate on a small-scale, in a pilot study, the effectiveness of experimental and industrial design.

### Significance

The study has helped establish the realization that any catalytic cycle contains common steps such as enzyme-substrate (ES) complex formation, activation of ES, transformation (or conversion) of complex reaction species (this includes the transition state species, TS, which either gets deactivated or proceeds to an enzyme-product (EP) complex), and dissociation of complex species such as ES and EP. The calculated life span of specific steps in the catalytic cycle presupposes that there are different first-order rate constants and consequently the possibility of different activation energies, both kinetic, the Arrhenius kind, and thermodynamic, the Gibbs kind. This should lead to a novel way of calculating activation parameters that may be of interest to all kinds of relevant engineers, technologists, and scientists in pure and applied sub-disciplines in industries and research institutes and institutions of higher learning.

## 5. CONCLUSION

All equations for the calculation of the life span of each step in the catalytic cycle (otherwise known as the reaction pathway) are derivable. There are always steps in an enzyme-catalyzed reaction in the catalytic cycle; the LS of each step is shorter than the total LS of deactivation and dissociation processes; this is applicable to forward reactions. The magnitude of the life span differs from one step to another, with the implication that there should always be different rate constants, and consequently, thermodynamic activation parameters must account for all FORC in accordance with the additivity principle. On matters of general interest, the Arrhenius and Eyring-Polanyi equations may, speculatively, be used for the determination of a fraction of the directly observable and measurable thermodynamic term, the enthalpy (thermal energy content of any system) by a graphical method. The values obtained can then be used to calculate the corresponding Arrhenius activation energy and free energy of activation, considering that the same plot is applicable to both kinds of activation energies. The fact that the sum of the different life spans is very much in agreement with the corresponding duration of either a backward or forward reaction is an indication of the validity of all the equations derived in this study.

## Acknowledgement

The management of the Royal Court Yard Hotel in Agbor, Delta State, Nigeria, is deeply appreciated for the supply of electricity during the preparation of the manuscript. The provider of the QuillBot grammar checker is thanked for the grammar in the manuscript.

